# Integrative analysis of *Hydra* head regeneration reveals actions of conserved signaling pathways and transposable elements

**DOI:** 10.1101/2022.05.17.492329

**Authors:** Aide Macias-Muñoz, Lisa Y. Mesrop, Heidi Yahan Liang, Ali Mortazavi

## Abstract

The extent to which animals can regenerate cells, tissues, or body parts varies largely. *Hydra* has a remarkable ability to undergo full body regeneration. Bisected polyps can regenerate the head and foot, and whole polyps can form from aggregates of cells. This capability is made possible by a cluster of cells known as the head organizer. Previous studies have found *Wnt3* and other developmental genes associated with head organizer function. Yet, the genetic and molecular mechanisms of regenerating *Hydra* heads remain unresolved. In this study, we used bulk RNA-seq, bulk ATAC-seq, and single-cell RNA-seq from a regeneration time course to characterize the gene expression, gene regulation, molecular, and cellular features of *Hydra* head regeneration. WGCNA modules and *cis*-regulation of candidate head organizer genes identified genes co-expressed with *Wnt3* and transcription factors important for regeneration. At the single-cell level, we identified at least six distinct cell types in the regenerating head tissue and characterized the expression of candidate genes. With these combined data sets, we identify and clarify some of the interactions between JNK, Wnt3, and EGRF signaling during *Hydra* head regeneration. Our study reveals coordination of early wound response, developmental transcription factors, and transposable elements during *Hydra* tissue regeneration and provides insight into the evolution of regeneration programs.

## Introduction

Tissue regeneration, which is the capacity to self-renew and differentiate into specific tissues, is a complex process involving the coordination of various cell types and genetic regulatory programs. The extent of regeneration in animals varies between organisms and tissue types, ranging from tissues with minimal regeneration capacities, such as mammalian heart and brain, to whole organism generation, such as in *Hydra* (Goldman & Poss 2020). *Hydra*’s capability of regenerating its body parts was discovered almost 300 years ago by Abraham Trembley (Trembley 1744). Following this discovery, additional studies have characterized the head and foot regeneration of *Hydra*. *Hydra* can fully regrow its head and tentacles 72 hours after bisection, ectopic heads can be induced from grafts, and adult polyps can be dissociated into cell aggregates capable of reforming a polyp (Browne 1909; Noda 1971; Technau et al. 2000; Bode 2003). This regrowth and patterning is made possible by a cluster of cells known as the head organizer that is responsible for axial patterning and head regeneration (Gierer et al. 1972; Browne 1909; Technau et al. 2000; Broun & Bode 2002; Bode 2012). While regeneration in *Hydra* is a long-standing topic of investigation, there are still gaps in knowledge regarding the temporal dynamics and the cellular and molecular basis of head regeneration. As a member of the Cnidaria phylum, which includes sea anemones, corals, and jellyfish, comparing the regeneration programs in *Hydra* to other cnidarians and to their sister group Bilateria will tells us about conservation and variations in gene function and gene regulatory network connectivity.

Previous *in situ* hybridization and chemical inhibition studies identified conserved developmental genes and transcription factors (TFs) associated with the *Hydra* head organizer. Some of the most well studied genes in *Hydra* are members of the Wnt family, especially *Wnt3*. *Wnt3*, *β-catenin* and *Tcf* genes are expressed in the hypostome of the adult *Hydra* polyp, the regenerating head, and the hypostome of a developing polyp formed by sexual budding (Hobmayer et al. 2000; Yukio Nakamura et al. 2011). *Wnt3* expression is controlled by two *cis*-regulatory elements that are directly influenced by Wnt/β-catenin signaling (Yukio Nakamura et al. 2011). Wnt genes *Wnt1*, *Wnt7*, *Wnt9/10a*, *Wnt9/10c*, *Wnt11* and *Wnt16* are also expressed in the adult hypostome, during budding, and during regeneration (Lengfeld et al. 2009). These genes are expressed in both epithelial germ layers (endoderm and ectoderm) of adult polyps, but their expression dynamics vary during budding and regeneration (Lengfeld et al. 2009). Homologs of the vertebrate axial patterning genes *Brachyury* and *Goosecoid* have also been associated with the head organizer. *Hydra Brachyury1* (*HyBra1*) is expressed in the endoderm of the adult hypostome, during budding, and early during regeneration (∼3 hours post bisection) (Technau & Bode 1999; Bielen et al. 2007). *HyBra2*, on the other hand, is expressed in the ectoderm layer of the hypostome and appears later during regeneration (between 6 to 15 hours) (Bielen et al. 2007). *Hydra Goosecoid* (*Gsc*) is expressed in the hypostome ectoderm and in the endoderm at the tentacle base of adult *Hydra* (Broun et al. 1999). During budding and regeneration, *Gsc* expression appears early in the endoderm, then is found in the ectoderm at the tip of the new hypostome and in the endoderm as a ring where tentacles will form (Broun et al. 1999). Notch, another conserved developmental pathway, is also implicated in the head organizer. Treatment of *Hydra* with DAPT, a Notch signaling inhibitor, results in malformed heads (Münder et al. 2013).

A traditional candidate gene approach provided strong evidence and validation for the actions of conserved signaling molecules and TFs. Yet, advances in sequencing technologies and generation of a *Hydra* genome then facilitated the identification of additional genes and TFs used in *Hydra* head regeneration (Chapman et al. 2010; Steele 2012). As an example, a proteomic and transcriptomic study characterized healing and patterning of the *Hydra* head (Petersen et al. 2015). The injury response and wound closure are associated with cell cycle control, nucleic acid binding, cytokinesis, cell signaling, and novel *Hydra* proteins (Petersen et al. 2015). Regeneration and patterning, on the other hand, are associated with developmental signaling pathways such as Wnt, TGF beta, Jak/STAT, MAPK and mTOR (Petersen et al. 2015). Another study found dynamic chromatin accessibility for transcription factor binding sites (TFBS) including paired box (Pax), forkhead box (Fox), SRY-related HMG-box (Sox) and Goosecoid (Gsc) during head regeneration (Murad et al. 2021). Moreover, a comparison of *Hydra* head and foot regeneration found that expression of some Wnt signaling genes including *Wnt3*, *Wnt9/10c*, *Wntless*, *Wnt7*, *dishevelled*, *sp5* and *β-catenin* is similar after injury and becomes head specific after 8 hours (Cazet et al. 2021). AP-1, Atf7, c/EBP, and Elf1 were identified as potential *Wnt3* regulators after injury (Cazet et al. 2021). These studies provide a suite of additional genes and TFs that may be involved in the actions of the head organizer.

Recent advances in single cell RNA-seq (scRNA-seq) technology and analysis software now allow for resolution at the single cell level. For model cnidarians, scRNA-seq has been used to describe cell types, genes specific to different cell types, and expression profiles of conserved developmental genes (Sebé-Pedrós et al. 2018; Siebert et al. 2019; Chari et al. 2021). In *Hydra*, scRNA-seq data from different developmental stages were clustered by cell linage (ie. endoderm, ectoderm, and interstitial) and cell types which included battery, stem, neuron, nematoblast, nematocyte, and gland cells (Siebert et al. 2019). In terms of regeneration, scRNA-seq was used to discover a newly described epidermis cell type that responds to injury and is responsible for wound healing in *Xenopus* (Aztekin et al. 2019). By contrast, scRNA-seq in axolotls uncovered that connective tissue cells with adult phenotype can revert to an embryo-like phenotype during regeneration (Gerber et al. 2018). These studies suggest that scRNA-seq can help us identify the cell types and molecular components involved in regeneration competency.

In this study, we used bulk RNA-seq and bulk ATAC-seq from a regeneration time course and hypostome to determine gene expression correlation and *cis*-regulation at the tissue level, respectively. We also used scRNA-seq from the adult *Hydra* hypostome and during a time course of regeneration to determine the cell types and genes involved in *Hydra* head regeneration. Using a weighted correlation network analysis (WGCNA) on bulk RNA-seq data, we identified a group of genes tightly co-expressed with *Wnt3* and genes important for injury response. We hypothesized that candidate regeneration genes would be co-expressed in regenerating tissue and in similar cell types. Using ATAC-seq data and pseudotime single cell trajectories for *Wnt3*-positive cell clusters, we identified candidate *Wnt3* regulators. Comparing the timing and potential functions of these genes reveals pathways that may be conserved in regenerating organisms. Lastly, using scRNA-seq, we identified at least 6 distinct cell types in regenerating tissues. Interestingly, head endoderm and ectoderm cells seem to undergo similar changes in expression of genes with uncharacterized functions and some associated with transposable elements. *Wnt3*, the proposed marker of the head organizer, development transcription factors, and candidate regeneration genes were predominantly expressed in the head epithelial cells. scRNA-seq served as a tool to validate previous findings and provide further evidence of gene function and interactions.

## Results

### Differential Expression and WGCNA of bulk RNA-seq

To identify signaling genes of the head organizer and candidate regulators of head regeneration, we did differential expression analyses and WGCNA on bulk RNA-seq data from a regeneration time course (Murad et al. 2021). Using differential expression analysis, we identified genes upregulated at t0 (immediately after head bisection) versus t12 (12 hours post-bisection) and t48 (48 hours post-bisection) (Figure S1). Genes upregulated in t0 had similar expression at 2, 4 and 6 hours post bisection (Figure S1). These are early injury response genes and include transcription factor genes *Otx1-like*, *cut-like*, *Meis-like*, *AP1*, *AP2*, *COUP*, *Myc-A-like* (Table S1). Based on expression trajectories, t12 encompasses a significant shift in gene expression during head regeneration and is when *Wnt3* peaks in expression (Murad et al. 2021). Along with *Wnt3*, genes upregulated at t12 include *Dkk1/2/4A*, *MAPK*, *Dl6x*, *Nkx*, *myc*, *Fos*, *Jun*, *Wnt2*, *Wntless* (Table S2). Genes upregulated at 48 hours include *HyBra1*, *HyBra2*, *HyAlx*, *Wnt1*, *Wnt8*, *Wnt9/10a* and *Wnt16* (Table S3).

Using WGCNA, we identified two modules of interest: coral2 module consisting of genes co-expressed with *Jun*/*Fos*, which are early wound response genes, and darkgreen module with genes co-expressed with *Wnt3,* the proposed *Hydra* head organizer gene (Srivastava 2021). These modules were chosen based having genes and expression trajectories of interest to healing and patterning. The coral2 module begins expression at 4 hours and peaks at 12 hours. This module increases early during the wound healing phase and is enriched for functions in cell growth and migration, epidermal growth factor response (EGFR) signaling, and immunity (Figure 1). Genes associated with this module included Toll (TLR) and tumor necrosis factor (TNFR) receptors (Table S4). Our findings support an immune response to injury in *Hydra* associated with early expression of *Jun*, *Fos*, TLR and TNFR signaling genes (Wenger et al. 2014).

**Figure 1.**
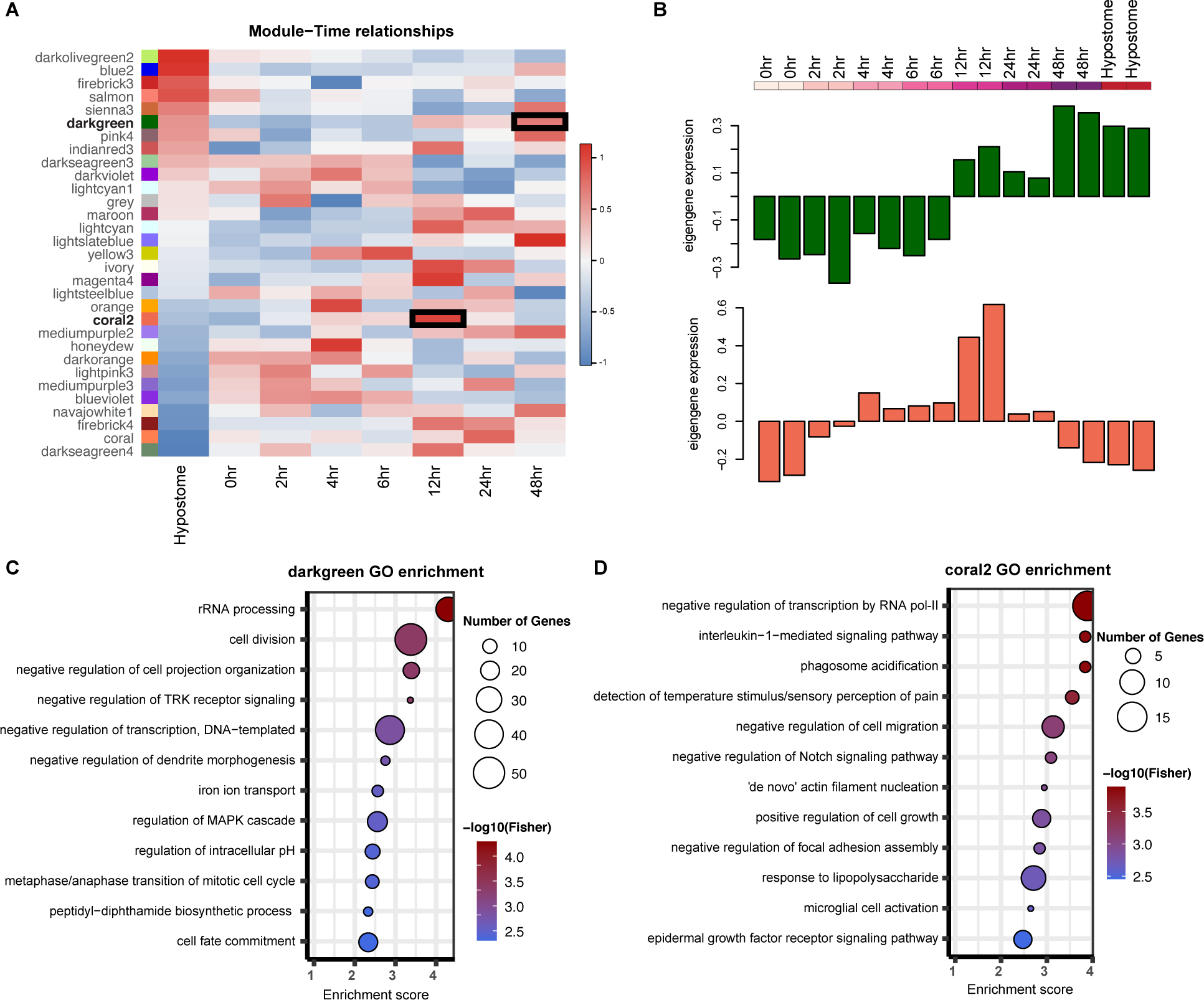
WGCNA reveals co-expression networks related to regeneration. A) Heatmap showing the correlations of module eigengenes to hypostome tissue and regeneration timepoints. *Wnt3* is found in the darkgreen module and Jun and Fos in the coral2 module. Boxes indicate significant correlations for our two modules of interest. B) Eigengene expression for the darkgreen and coral2 modules during head regeneration. C) Gene Ontology (GO) enrichment for the darkgreen module > 5 genes. Size of circle indicates number of genes in the GO term category and color is the statistically significant p-value score in logarithmic scale. D) GO enrichment for the coral2 module.

Unlike in coral2, the darkgreen module is associated with increased expression later during regeneration at t12, peaking at t48, and being highly expressed in the adult hypostome (Figure 1). Wnt signaling is highly conserved in evolution and is known to play a role in cell proliferation, cell fate determination, cell migration during development and regeneration. As expected, the co-expression module containing *Wnt3* is primarily involved in cell division, cell fate commitment, and differentiation most likely directing cell differentiation and growth during regeneration. Genes controlling cell cycle and cell death are known to be expressed during regeneration and are enriched in this co-expression module, such as cyclin-dependent kinases (CDKs), receptor type protein tyrosine phosphatases (RPTPs), and ATM serine kinases (Petersen et al. 2015). While the coral2 genes are responding to injury and involved in wound healing, the genes in the darkgreen module are active in regeneration and patterning. Some of the genes in this module include Wnt signaling such as *Wnt7*, *Wnt9/10a*, *Wnt9/10c*, *Dishevelled*, and *Sp5* (Table S5). Other transcription factors include *COUP*, *Erg-like*, *Meis1-like*, *SoxB2-like*, *SoxE1-like*, *Rax-like*, *PaxC*, *TEF1*, *E3*, *FoxJ1*, *DMRT3*, and *SRY* (Table S5). Of note, this cluster also has EGF-domain containing genes, mobile elements, and genes associated with reverse transcription.

### Candidate head organizer gene cis-regulation

Previous studies using ATAC-seq and ChIP-seq identified peaks at the *Hydra Wnt3* promoter and one upstream putative-enhancer region (Y. Nakamura et al. 2011; Cazet et al. 2021; Murad et al. 2021). In a wound response study, transcription factor binding motifs (TFBM) Atf7 and Isl1 were found in the *Wnt3* promoter (Cazet et al. 2021). To identify which genes may be signals of the head organizer, we searched the *cis*-regulatory regions of canonical Wnt3 signaling and injury response genes for footprint signatures of bound transcription factors. Using combined hypostome ATAC-seq data (Siebert et al. 2019; Murad et al. 2021), we identified TFBS motifs that are common across some Wnt signaling and early wound response genes (Figure 2; Table S6). TFs involved in wound response include CREB, Jun, and Fos. We found matches for CREB in the peaks adjacent to *Wnt3*, *Dishevelled*, *TCF*, and *Wnt7* and *Wntless* (Figure 2; Table S6). We found matches for Jun and Fos in peaks near *CREB*, *Dishevelled*, *Dkk1/2/4A*, *Jun*, *Sp5*, *TCF*, *Wnt7*, and *Wntless* (Figure 2; Table S6). The TFBS motifs that were overrepresented in our genes of interest included Iroquois complex factors ara/caup and mirror, Arid3a, Sox, and optix. Interestingly, the other group of TFBS motif that appeared together were Jun, Fos, CREB and Atf. These *cis*-regulatory regions are potentially of important to injury response and healing. The last group of TFBS motifs that appeared together were onecut, Dux, and NF-Y. The areas that have these motifs are likely important for patterning. Lastly, of our genes of interest, the *Wnt3* and *TCF* promoters were the most similar in their *cis*-regulatory regions.

**Figure 2.**
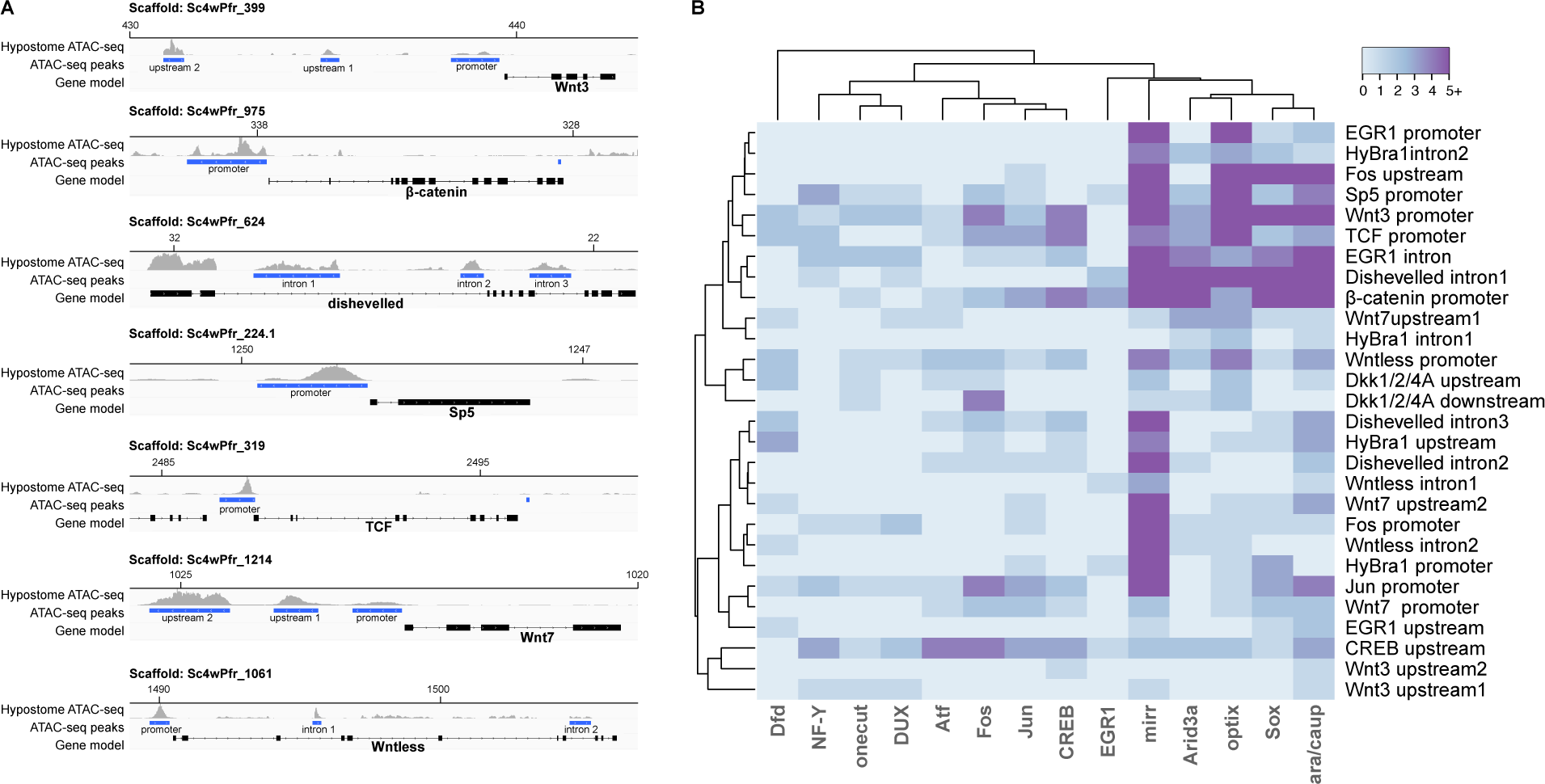
*Cis*-regulation of injury response and Wnt signaling genes. A) Diagram of Wnt signaling genes showing the gene region, ATAC-seq peaks called, and ATAC-seq read coverage. B) Diagram of TFBS motifs present in peaks adjacent to candidate genes.

### *C*ell types of the regenerating Hydra head

Single cell RNA was collected from the *Hydra* adult hypostome and at three time points during head regeneration: 0, 4, and 12 hours post bisection (Figure 3). Single cell RNA-seq libraries were sequenced to an average depth of ∼80 million reads per library (Table S7). We recovered ∼500-3,000 cells per sample after filtering (Table S7; Figure S2) for a total of 8626 cells. The cells grouped into 13 clusters (Figure S2). These clusters were defined by the most highly differentially expressed genes, which we refer to as top markers. We determined the identity of these 13 clusters based on the functions of top markers (Table S8), their expression in the *Hydra* single cell atlas available at https://singlecell.broadinstitute.org/, and expression of known *Hydra* cell markers (Figure S3) (Siebert et al. 2019). Based on these candidates for different cell types, we confidently identified six distinct cell types in the *Hydra* regenerating head: head epithelial ectoderm (1444 cells), interstitial stem (169 cells), neuron (271), nematoblast (164 cells), and nematocyte (170 cells). We also found clustering of cells with both endoderm and ectoderm head epithelial markers (head epithelial 1-3526 cells, head epithelial 2-2006 cells, and head epithelial 3-255 cells), and 3 clusters of unidentified cells (unidentified 1-61 cells, unidentified 2-473 cells, and unidentified 3-87 cells).

**Figure 3.**
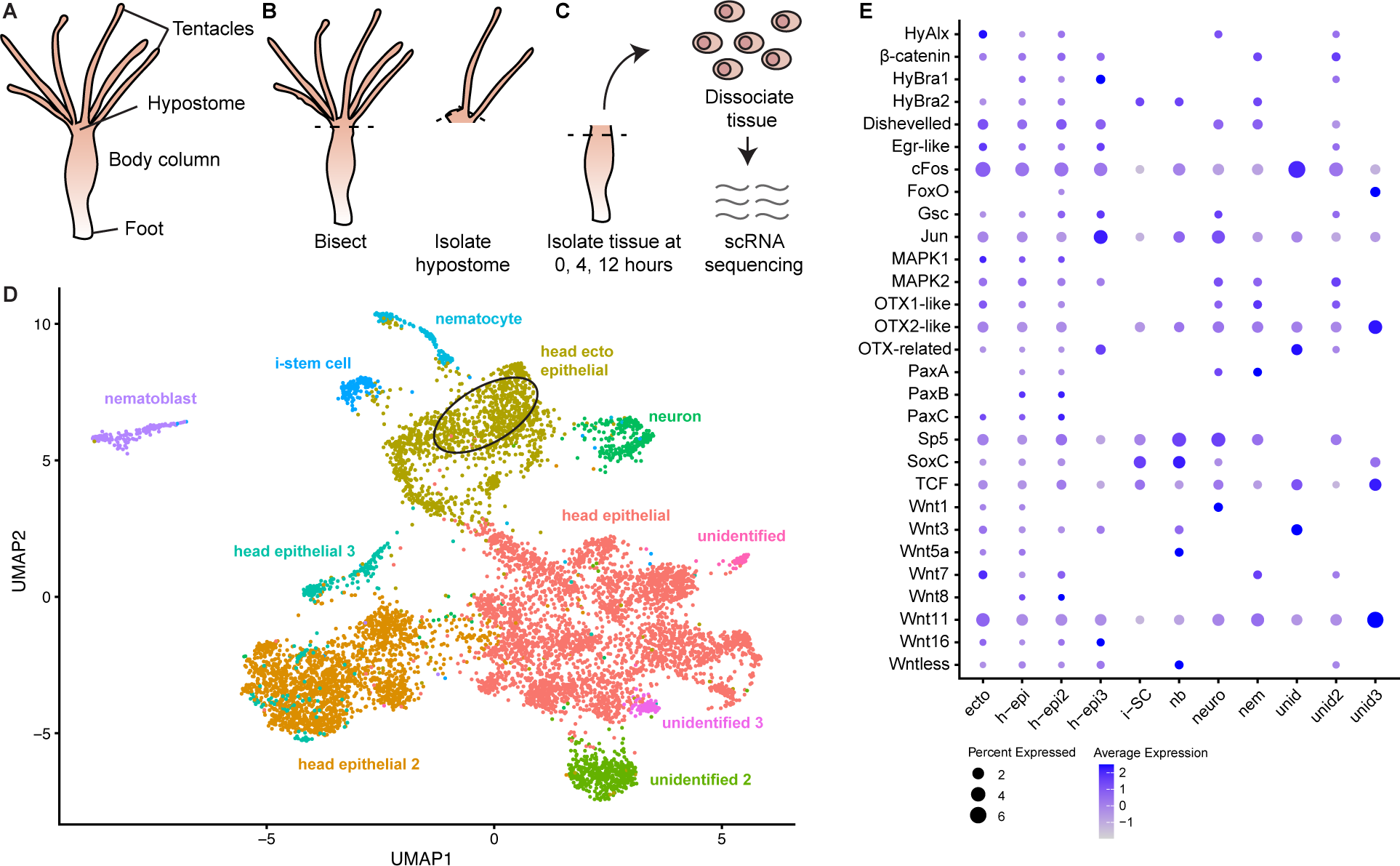
Cell clustering and candidate gene expression of regenerating *Hydra* heads. A) The *Hydra* body plan consists of tentacles, hypostome, body column, and foot. Circle denotes an area where some *Wnt3*-positive cells are clustered. B) For the hypostome sample, *Hydra* were bisected below the tentacle ring, the tentacles were removed, and the remaining tissue was collected. C) For a regeneration time course, *Hydra* were bisected below the hypostome, and the adjacent tissue was collected at 0 hr, 4 hr, and 12 hrs. The dissected tissues were dissociated into single cells, mRNA was extracted and sequenced. D) UMAP of cell clusters annotated with cell types. E) Expression of candidate regeneration genes in the depicted cell types.

### Expression of candidate regeneration genes

Previous RNA-seq studies identified genes involved in *Hydra* head regeneration (Petersen et al. 2015; Cazet et al. 2021; Murad et al. 2021). We looked at the distribution and expression of these candidate regeneration genes in our scRNA-seq clusters (Figure 3E). *Wnt3* is the most well studied gene involved in *Hydra* head regeneration. We found *Wnt3* was predominantly expressed in the head ectoderm cluster (Figure S3). Members of the Wnt signaling pathway that increase in expression early as a response to injury and become hypostome specific after 8 hours include *Wnt3*, *Wnt7*, *Wnt9/10c*, *Wntless*, *β-catenin*, *dishevelled* and *Sp5* (Cazet et al. 2021). We hypothesized that these genes would be expressed in similar cells or cell clusters to *Wnt3*. While the expression and distribution of these genes varied, all genes except *Wnt9/10c* were expressed in the head ectoderm and epithelial clusters (Figure 3E). In addition to *Wnt3*, *Jun*, *Fos*, and *HyBra1* were found to have dynamic spatial and temporal expression in *Hydra* (Petersen et al. 2015; Cazet et al. 2021; Murad et al. 2021). *Jun* and *Fos* are hypothesized to be conserved early wound response genes in regeneration (Morello et al. 1990; Kamp et al. 1995; Wenemoser et al. 2012; Sabin et al. 2019; Srivastava 2021). In our dataset, both *Jun* and *Fos* were highly expressed in the head epithelial cells (Figure 3E). Similar to *Jun* and *Fos*, *Brachyury* is also proposed to have a conserved role in animal patterning (Tewari et al. 2019). In *Hydra*, *HyBra1* functions in head formation with expression localized to the hypostome in adults and developing buds (Technau & Bode 1999). *HyBra1* is mostly in the endoderm while its paralog *HyBra2* is predominantly expressed in the ectoderm (Bielen et al. 2007). In our single cell analysis, *HyBra1* was expressed in the three head epithelial clusters and *HyBra2* was the paralog expressed in the ectoderm cell cluster (Figure 3E). Overall, development and candidate regeneration genes were found expressed in head epithelial cells during head regeneration (Figure 3E). This result is consistent with *Hydra* lacking interstitial cells maintaining the ability to undergo regeneration (Marcum & Campbell 1978). Looking specifically at *Wnt3*-positive cells, we found *Wnt16*, *Otx2-like*, *Egr-like*, *HyBra2*, *Jun* and *Fos* co-expressed with *Wnt3*. These genes are known to be involved in regeneration and early wound response in other animals (Petersen et al. 2015; Srivastava 2021). The co-expression of these genes in individual cells suggests a role for these pathways in *Hydra* head organization. This result supports results from WGCNA.

To investigate gene expression of regenerating epithelial tissue in pseudotime, we extracted cell clusters excluding nematocyts, nematoblast, neuron and stem cells. This analysis revealed a trajectory where *Wnt3* expressing hypostome cells appeared late in pseudotime (Figure 4A). To visualize the expression of candidate genes we mapped Wnt signaling genes, injury response, and candidate regulators in pseudotime (Figure 4; Figure S4). As expected, we observed *Wnt3* to have high expression early, followed by a decrease and then increase in expression (Figure 4B). A slingshot analysis supported these results (Figure S5). Some of the genes that had similar expression trajectories included *β-catenin*, *HyBra1*, *Jun*, *EGR-like*, *TCF*, *SoxC*, and *Sox18*. *Sp5* which is a *Wnt3* agonist has the opposite trajectory to *Wnt3*, potentially increasing in expression as an antagonist regulator after injury response. Of the transcription factors whose binding motifs are overrepresented in our genes of interest, *Irx* which binds ara/caup and mirr has similar expression to *Wnt3*. Genes matching *homeodomain (Dux)* and *NF-Y* have very high expression early in pseudotime that drops to a baseline expression.

**Figure 4.**
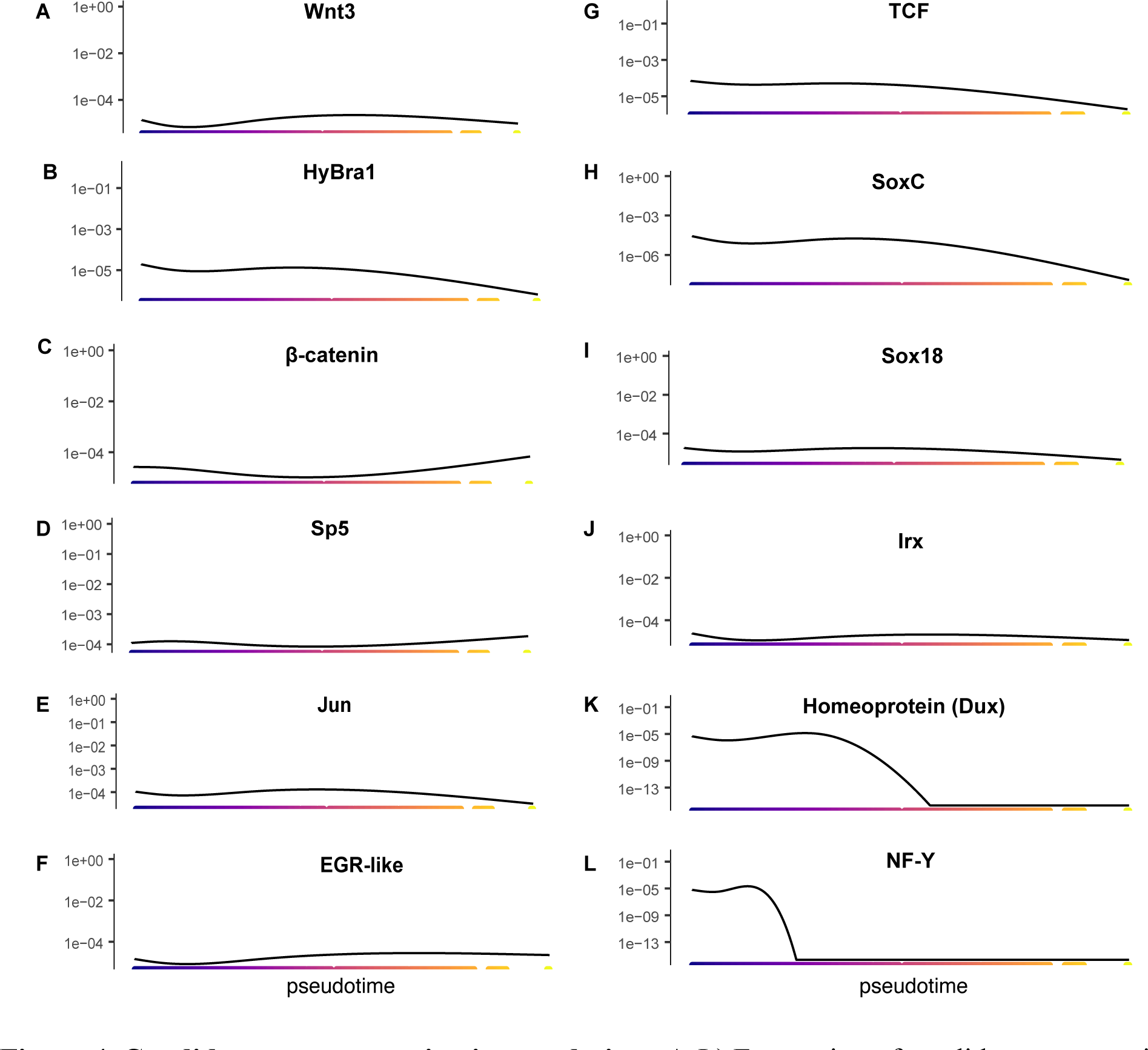
Candidate gene expression in pseudotime. A-L) Expression of candidate regeneration genes and transcription factors in increasing pseudotime. Y-axis scale shows early regeneration pseudotime in purple and late regeneration in yellow.

## Discussion

The cellular organization and actions of the *Hydra* head organizer are topics of high interest in comparative developmental biology. While there have been fundamental studies investigating candidate genes using *in situ* hybridization and chemical inhibition, we can now expand on these findings using high throughput sequencing to identify additional genes and pathways that allow for head regeneration. In this study, by re-analyzing bulk RNA-seq and ATAC-seq data, we confirm the role of conserved injury response and patterning pathways and identify transcription factors that may play a role in regulating *Hydra* head regeneration. We have generated a cell atlas and cell trajectories for *Hydra* head regeneration.

Although we did not identify the same cell diversity found in whole *Hydra* (Siebert et al. 2019), we saw distinct separation of stem, neuron, nematoblast, and nematocyte cells. The clustering of head epithelial cells was driven by genes associated with uncharacterized proteins, ribosome binding, zinc metalloproteinase, and reverse transcriptase (Table S8). These top markers occluded the biological differences of endoderm and ectoderm cells. Results imply that there are similar gene expression profiles in regenerating epithelial tissues. While endoderm and ectoderm are distinct cell types, injury drives the convergent expression of some genes including reverse transcriptases and mobile elements. This brings about a question as to the role of transposable elements in regeneration.

### A role for transposable elements during Hydra head regeneration

Approximately 50% of the *Hydra* genome is made up of transposable elements (Chapman et al. 2010). The organismal function of transposable elements (TEs) remains largely unknown with some hypothesizing that they function in gene regulation (Slotkin & Martienssen 2007; Bourque 2009). In mammals, TEs are highly expressed in embryos during zygotic genome activation and in pleuripotent cells at different stages (Gerdes et al. 2016; Hackett et al. 2017; Low et al. 2021). TEs have been shown to affect chromatin accessibility and even to have post-transcriptional effects thus it is hypothesized that they have a role in pleuripotency (Hackett et al. 2017). Whether TE expression is due to global opening of the chromatin or whether TEs play a direct role in regulating shifts in expression is still being investigated (Gerdes et al. 2016; Low et al. 2021). In terms of regeneration, increase in TEs were detected in regenerating haematopoietic stem cells and overexpression of TEs led to their activation (Clapes et al. 2021). In the sea cucumber *Holothuria glaberrima*, it was found that changes in transcription of retrotransposons were associated with tissue regeneration (Mashanov et al. 2012). Moreover, it was recently hypothesized that proper regulation of TE activity is important for regeneration competency across animals (Angileri et al. 2022).

In our study, several genes related to reverse transcription and transposable elements were top markers for the single cell data and influenced clustering (Table S8). Removal of these genes in the clustering analysis may have clarified the distinction between epithelial cells, but that would have resulted in missing an interesting signal of TE expression changes. Previously it was found that in *Hydra*, retrotransposons increase in expression from 0 to 12 hours during head regeneration (Petersen et al. 2015). This trend was recapitulated in the bulk RNA-seq data, where we saw peak expression of some TEs at 12 hours post bisection that correlated with peak of expression in *Wnt3* (Table S5). To identify the TEs that may have a significant role in regeneration, we searched Dfam database for top markers of the single cell data or those co-expressed with *Wnt3* (Storer et al. 2021). TEs that had matches included transib (Class 2), L2 (CR1), Zenon1 (CR1) and Zenon2 (CR1). These findings suggest a role for TEs in *Hydra* head regeneration particularly CR1/L2 LINE elements at 12 hours post bisection. While not a lot of research has been done to investigate the function of TEs in *Hydra*, one study found expansion of these LINE elements associated with a larger genome in *Hydra vulgaris* and close relatives (Wong et al. 2019). Future work should focus on characterizing the identity of these uncharacterized TEs and determining their function. Specifically, it is of interest to investigate where TEs have a direct role in regulating the expression of candidate regeneration genes.

### WNT, MAPK, and EGFR signaling important for Hydra regeneration

The actions of canonical *Wnt3* signaling on *Hydra* head regeneration have long been known. Previous studies found similar trajectories of expression in some Wnt3 signaling genes including *Wnt3*, *Wntless*, *Wnt7*, *dishevelled*, *sp5* and *β-catenin* (Cazet et al. 2021). We found these genes expressed in epithelial cells and we found their expression increasing or peaking after injury in a pseudotime analysis. Tracking their expression using bulk RNA-seq, confirms that these genes are co-expressed with *Wnt3*. These known canonical Wnt signaling genes had similar *cis-* regulation by transcription factors including ara, CEBP, CREB, ara, and optix. In addition to these Wnt signaling genes, we found the *Wnt3* antagonist *Dkk1/2/4A* to peak around t12. While in intact *Hydra* this gene is expressed in the body column below the hypostome, after injury it increases in expression at the site of amputation (Guder et al. 2006). In this study, we find the expression of this gene correlated with *Wnt3*, *Jun* and *Fos* all peaking at 12 hours post head bisection. Interestingly, we found that *Dkk1/2/4A* has TFBS motif for Jun and Fos similar to *Wnt7*, *dishevelled*, and *sp5*. A previous study suggested that Jun could be the molecule to triggers *Dkk1/2/4A* for tissue repair (Guder et al. 2006). Here, we show that it likely does so through *cis*-regulation. These results demonstrate that multiple mechanisms of *Wnt3* agonists and antagonists are at play during regeneration.

*Jun* and *Fos* are part of the JNK (c-Jun N-terminal kinase) pathway which is one of the MAPK (mitogen-activated protein kinase) signaling pathways responsible for cell proliferation and apoptosis. The JNK signaling pathway is known to function in early wound response. In *Hydra*, the MAPK signaling pathway is activated following injury, cells then undergo apoptosis and release *Wnt3*. Thus MAPK and specifically JNK are necessary to trigger *Wnt3* signaling for *Hydra* regeneration (Chera et al. 2011; Tursch et al. 2022). In this study, we see *Jun* and *Fos* expressed in all cell types of the regenerating *Hydra* head, and *MAPK1* and *MAPK2* in the epithelial cells. In pseudotime, these genes peak in expression following injury. Bulk RNA-seq reveals that they begin increasing in expression earlier than *Wnt3,* but they peak with *Wnt3* at 12 hours post head bisection. *Jun* and *Fos* were co-expressed with *Wnt3* at the single cell level. While not directly regulating *Wnt3*, *Jun* and *Fos* are directly affecting genes downstream and upstream of Wnt3 signaling. Some transcription factors common to *Jun*, *Fos* and *Wnt3* include ara, caup, CEBP, Dux, onecut, optix, and Sox. These results suggest a role for these transcription factors upstream of signaling pathways for injury response and patterning.

Moreover, while the functions of EGFR in regenerating *Hydra* have not been established, in *Drosphila*, EGFR and MAPK act downstream of JNK signaling during apoptosis-induced proliferation (Fan et al. 2014). In the planaria *Schmidtea mediterranea*, silencing EGFR genes results in malformation during regeneration due to effects on cell proliferation and differentiation (Fraguas et al. 2011). In this study, we identify EGF-containing domains as top markers of regenerating hypostome and enrichment of EGFR signaling in the module where *Jun*, *Fos*, and *MAPK* are grouped. These combined results uncover a potential role for EGFR signaling during *Hydra* head regeneration.

### Evolution of regeneration pathways

To understand the degree to which regeneration is conserved or divergent across species, we need to compare the underlying genes and regulatory programs in regenerating species of different taxa. A comparison of genes involved in head and tail regeneration between planaria and *Nematostella* is summarized by Schaffer et al. 2016 (Schaffer et al. 2016). This study found orthologous genes expressed in anticipated patterns (based on the organisms oral and aboral development) for axial regeneration such as *Otx* and *Six* (Schaffer et al. 2016). We find both genes associated with head regeneration in *Hydra*. Genes that were associated with head regeneration in both *Nematostella* and planaria included *SoxB*, *Wnt2*, and *FoxD* (Schaffer et al. 2016). In *Hydra*, members of these gene families were expressed in epithelial cells, but we are unable to state whether these are paralogs or orthologs that have a conserved function in head regeneration. A similar pattern arises when we look at Wnt genes in *Hydra* and the acoel *Hofstenia*. In *Hofstenia*, *Wnt3* and other genes associated with *Hydra* head regeneration were found to function in posterior regeneration. These genes included *Brachyury*, *Sp5* and *FoxA1* (Ramirez et al. 2020). Moreover, Egr is the proposed regeneration initiation factor in *Hofstenia.* In a comparison of chromatin accessibility during regeneration between *Hostenia* and a planaria Egr was one of the most accessible motifs along with Fox and Jun/Fos. Here we found *Egr-like*, *Jun*, and *Fos* expressed co-expressed with *Wnt3*. This comparison across species highlights the roles of Wnt signaling and TF activity of Egr, Wnt, Fox, Jun, Fos, Sox, Otx and Six in regeneration and patterning.

The actions of canonical Wnt signaling have been explored and established as a pathway important to regeneration not only in cnidarians, planaria, and acoels, but also in sea stars, axolotls, *Xenopus*, zebrafish, and even mice (Macias-Muñoz in prep). The role of this pathway in regeneration across organisms suggests it is either deeply conserved or is crucial to patterning and is constantly being co-opted. Additional comparative studies in regenerating organisms should determine the regulatory networks of Wnt signaling genes (Srivastava 2021), and also whether the signaling cascades are exactly the same across organisms. This will reveal the extent of conservation. Similarly, the actions of MAPK as wound response triggering Wnt for repatterning seem to be found across regenerating animals and in different tissues (Zhang et al. 2014). In this study we find that Jun and Fos have TFBS near Wnt3 signaling components and antagonists. Whether MAPK and/or Wnt signaling directly interacts with EGFR during regeneration is an open question. In this study we find signatures of EGFR signaling during wound response and early regeneration. There is increasing evidence of potential crosstalk between EGFR with JNK and Wnt signaling in development and cancer (Paul et al. 2013; Kushnir et al. 2017). Future research should investigate whether crosstalk occurs in animal regeneration.

The crosstalk between signaling pathways, especially the conservation of interactions is important in determining the features of healing and patterning that are important for regeneration competency. Also important at the molecular level, is identifying the transcription factors that play a role in regulating these signaling cascades during regeneration. An overview of TFBS for genes important to *Hydra* regeneration revealed potential interactions through *cis*-regulatory mechanism (Figure 2). In addition, this line of inquiry revealed potential actions of TFs such as ara, mirr, caup, sox and optix. Araucan (ara), caupolican (caup) and mirror (mirr) are three closely related homeobox transcription factors that make up the Iroquois complex in *Drosophila*. These genes are responsible for prepatterning in *Drosophila* and vertebrates where they have undergone expansions (6 in humans) (Kim et al. 2012). Here we find peaks and potential binding of these factors near many of the genes that are important for regeneration. Irx also has a similar expression trajectory in pseuditme to the *Wnt3* head organizer genes. This finding suggests a role for Iroquois homeobox genes as important for tissue patterning during regeneration.

Transcription factors sox and six (optix) may also be important for patterning in regeneration and may have conserved functions across organisms. Multiple Sox genes have been shown to have regulatory functions for Wnt signaling. As an example, Sox2 interacts with Wnt signaling and is important for neuron regeneration in sea star larvae (Kormish et al. 2010; Zheng et al. 2022). Sox13 is a TCF repressor and is proposed to control Wnt signaling in embryo development (Marfil et al. 2010). In *Xenopus*, XSox17α, XSox17β, and XSox3 bind β-catenin to inhibit Wnt signaling during embryo development (Zorn et al. 1999). Recent work suggests a link between *Sox18* and Wnt signaling in cancer (Kormish et al. 2010; Yin et al. 2017; Geng et al. 2020). As Sox has been shown to directly modulate Wnt signaling in vertebrates, we predict that one or more of the *Sox* genes play a role in *Wnt3* signaling in *Hydra*. One question of interest is whether the functions of Sox as a Wnt regulator are conserved or if these actions have been co-opted in animal evolution. In the examples above Sox repress Wnt by two different mechanisms, binding β-catenin or TCF. Our data suggests candidate Sox are acting directly as Wnt agonists in *Hydra* regeneration by binding a *cis*-regulatory site. These comparisons suggest co-option of different Sox family members at different points in the signaling cascade, but a constraint on the evolution of this gene regulatory network (Kopp 2009).

*Optix* is a member of the sine oculis homeobox (SIX) gene family. *Six3/6* functions in patterning in *Nematostella* (Sinigaglia et al. 2013). Members of the *Six* gene family have been found to have roles in eye regeneration in jellyfish and newts (Stierwald et al. 2004; Grogg et al. 2005). In our analyses, we found a bound TFBS for Optix (Six3/6) at the *Wnt3* promoter. These results indicate that *Six3/6* could also be a *Wnt* regulator. In mice, Six3 and Six6 repress Wnt signaling (Diacou et al. 2018). It could be that members of the *Sox* and *Six* gene family have a role in Wnt signaling across species. In this case it is the homologous *Six3/6* in multiple species, but the direct regulation of Six TFs on *Wnt* or their mechanisms of interaction have not been functionally described.

Overall results from our study and previous studies highlight that some of the same gene families important to development are used by animals for regeneration. However, the expression trajectories and genetic programs underlying patterning vary between normal development and regeneration. Moreover, while the same gene families might be used, the gene family members used and their expression in oral versus aboral regeneration can vary across organisms. Instances where gene network connections appear conserved, such as the role of Sox and Six in *Wnt* signaling regulation, is where future work should focus. Similarly, TEs seem to play a role in regeneration competency and pluripotency in different taxa. While the direct role of TEs is yet unknown, here we find that an increase in TE expression is correlated with shifts in gene expression suggesting a direct role in gene regulation. Investigating how the cellular and genetic underpinnings of regeneration are similar across animals, can lead to insights about the evolutionary history of regeneration and the features that underlie regeneration competency.

## Conclusions

The cellular organization and actions of the *Hydra* head organizer are topics of high interest in comparative developmental biology. While there have been fundamental studies investigating candidate genes using *in situ* hybridization and chemical inhibition, we can now expand on these findings using single cell transcriptomics to identify additional genes and pathways that allow for head regeneration. In this study we have generated a cell atlas and cell trajectories for *Hydra* head regeneration. We were able to identify at least 6 cell types in the regenerating tissues. In addition, by generating cell trajectories for regenerating tissues we identified potential regulators of Wnt/β-catenin signaling including members of the Six and Sox gene families. By re-analyzing bulk RNA-seq and ATAC-seq data, we find that co-expression and binding patterns that are confirmed by our single cell data, further supporting our findings.

## Materials and Methods

### Animal care

*Hydra vulgaris* (strain 105) were maintained in glass pans in the laboratory at room temperature in *Hydra* medium (4.22 g calcium chloride dehydrate, 3.06 g magnesium sulfate anhydrous, 4.2 g sodium bicarbonate, 1.1 g potassium carbonate in 100 liters of DI water). *Hydra* were fed brine shrimp (*Artemia salina*) twice per week and cleaned after every feeding. To clean the containers, *Hydra* medium with brine shrimp was poured out and any detached *H. vulgaris* were recovered. New *Hydra* medium was added so that fed *H. vulgaris* were left submerged in about 2 inches of medium. *H. vulgaris* were starved 1-3 days prior to RNA extractions to avoid brine shrimp contaminations.

### Differential Expression Analysis of bulk RNA-seq

Smooth quantile normalized read count data were obtained from Murad et al. for timepoints 0, 2, 4, 12, 24 and 48 hr after head bisection (Murad et al. 2021). Data were analyzed using edgeR v. 3.40.2. MDS plot revealed similarities early in regeneration (t0-t6) and at timepoints t12 and t24. Due to this clustering, we did pairwise differential expression analyses using glmQLFTest for t0 and t12, t0 and t48, and t12 and t48 to identify genes upregulated (P-value > 0.5) at early, mid, and late regeneration.

### WGCNA analysis on bulk RNA-seq

Bulk RNA-seq data of the *Hydra* hypostome tissue and head regeneration time points (0, 2, 4, 6, 12, 24, and 48 hours) were obtained from GEO accession GSE127279 (Murad et al. 2021). The dataset consists of a total of 16 samples, with 2 biological replicates for both the hypostome tissue and for each head regeneration time point. Raw reads were mapped to the reference transcriptome (Murad et al. 2021) using Kallisto (Bray et al. 2016).

Weighted gene co-expression analysis was performed using the WGCNA package v. 1.70-3 in R (Langfelder & Horvath 2008). Genes with counts less than 10 in more than 2 samples were removed to reduce noisy, non-significant correlations. After filtration of lowly expressed genes, the input expression matrix consisted of 16,971 genes. Expression of these genes was normalized with the variance-stabilizing transformation function of DESeq2 v. 1.28.1 (Love et al. 2014). To achieve a network with scale-free topology, a soft threshold of 18 was selected using the “pickSoftThreshold” in WGCNA package and the adjacency matrix was created with a “signed” network. Modules, or groups of genes that are highly co-expressed across samples were generated using a hierarchical clustering algorithm. A threshold of 0.2 and minimum module size of =30 were used to merge similar expression profiles to obtain 31 modules. The *Wnt3* co-expression network has over 1000 genes and because visualization of all the genes and their interactions would be difficult to illustrate, we only selected genes the top 38 annotated genes that have strong connectivity with *Wnt3*.

The *Hydra* transcriptome was annotated using Blast2GO (Conesa et al. 2005). GO terms enrichment analysis for candidate modules was carried out using TopGO v. 2.41.0 (Alexa & Rahnenfuhrer). GO enrichments of the module containing *Wnt3* and all the co-expressed genes were used as the test dataset and the GO enrichments of all the annotated genes from the transcriptome were used as the universal dataset. To determine overlap between the *Wnt3* co-expression network generated from the RNA-seq data of the hypostome tissue and head regeneration timepoints and Seurat clusters, we performed a Fisher exact test to determine the significance of the overlap.

### Identification of Wnt3 Transcription Factor Binding

Raw read ATAC-seq data for the *Hydra* hypostome were obtained from GEO GSE127277 (Murad et al. 2021) and GEO GSE121617 (Siebert et al. 2019). Reads were trimmed using Trimmomatic v. 0.39 (Bolger et al. 2014). After trimming, reads were mapped to the *Hydra* mitochondrial DNA and genome using bowtie2 v. 2.4.1 (Langmead & Salzberg 2012). Mitochondrial reads were removed using Picard v. 2.26.3. Homer v. 4.11.1 was used to call peaks. Peaks and bam files for the five biological replicates were then merged before performing downstream analysis with TOBIAS v. 0.12.10 (Bentsen et al. 2020).

### Single cell regeneration time course and cell dissociation

12-15 *H. vulgaris* individuals were used for each sample time point for a total of 3 timepoints (t0, t4 and t12) and the hypostome (Figure 1B). For the hypostome sample, the head was bisected below where the tentacles attach, the tentacles were removed, and the remaining tissue was collected (Figure 1B). For t0, the head was bisected, and the tissue directly below was immediately collected (Figure 1B). For timepoints t4 and t12, the hypostome was bisected then *Hydra* were left in medium at room temperature to heal and regenerate for 4 and 12 hours, respectively. For each time point, the regenerated tissue was collected (Figure 1B). The collected tissues were pooled in a 1.5 ml tube for each sample and washed with *Hydra* medium. The medium was then removed and *H. vulagris* were dissociated using Pronase E as previously described (Greber et al. 1992). Briefly, 1 ml of *Hydra* dissociation medium (5 mM CaCl_2_ 2H_2_O, 1 mM MgSO_4_ 7H_2_O, 2.8 mM KCL, 11 mM TES, 0.67 mM Na_2_HPO_4_, 0.44 mM KH_2_PO_4_, 5 mM Na Pyruvate, 5 mM Na_3_ citrate 2H_2_O) with ∼75u/mL Pronase E (VWR, Radnor, PA) was added to each sample. Samples were placed on a nutator for 90 minutes. Cells were washed through a 70µm cell strainer (Corning, Corning, NY) then centrifuged at 300xg for 5 minutes at room temperature. Pellets were resuspended and strained through a 40µm cell strainer (Corning, Corning, NY). The final sample was resuspended with 60 µl 1X PBS containing 0.01% BSA. Cells were counted on a TC20 Automated Cell Counter (Bio-Rad, Hercules, CA).

### scRNA sequencing

Immediately after dissociation, cells were diluted to 2500 cells/µl in 1X PBS containing 0.1% BSA. 4.5 µl of sample were used to generate single cell RNA-Seq libraries by following the Illumina Bio-Rad SureCell WTA 3’ Library Prep Guide (Illumina, San Diego CA and Bio-Rad, Hercules, CA) and using the ddSEQ Isolator for Single Cell Sequencing (Bio-Rad, Hercules, California). Libraries were quantified and quality checked using a bioanalyzer (Agilent, Santa Clara, CA). Libraries were then multiplexed and sequenced using the setting “paired end, R1:68 cycles, R2:75 cycles” on a NextSeq 500 sequencer (Illumina, San Diego, CA).

### scRNA-seq clustering and pseudotime

Individual cells were identified from demultiplexed libraries using ddSeeker v.1.2.0 (Romagnoli et al. 2018). A transcriptome from Murad et al. (Murad et al. 2021) was indexed using kallisto v. 0.46.1 (Bray et al. 2016). Reads were mapped to the transcriptome according to kallisto using default settings. Abundances were grouped into a matrix for each sample using a custom script. Counts were then collapsed to gene level by adding transcripts matching the same gene ID of the *Hydra* 2.0 genome (Chapman et al. 2010). Counts data were loaded into R and analyzed using Seurat v. 4.3.0 (Butler et al. 2018; Stuart et al. 2019). Briefly, Seurat objects for each sample were made using min.cells = 1, min.features = 120. Each object was then assigned a time during regeneration (t0, t2, t4, t12, hypo). The 5 individual objects were then merged into one large Seurat object. The Seurat object was normalized using sctransform v. 0.3.5 and harmony v. 0.1.1 to remove batch effects. Cells were then clustered and visualized using UMAP.

For pseudotime analysis, we extracted gene expression information for the epithelial clusters. Expression data were input into monocle3 v. 1.2.7 (Qiu et al. 2017). After roviding Seurat clustering results, we constructed single-cell trajectories and visualized all clustered cells in pseudotime. Candidate genes were visualized for expression in pseudotime along branches of interest. Similarly, to confirm results, we input data from Seurat into slingshot v. 2.10.0. We input clustering from harmony and calculated lineages and pseudotime. To determine if *Wnt3* expression increased late in pseudotime, as would be expected for the reestablishment of a head organizer, we visualized expression of *Wnt3* in slingshot pseudotime.

## Data and code availability

Raw single cell RNA-seq data and counts matrices with counts generated and analyzed for this paper are available under GEO accession GSE193277. R and shell scripts accompanying this paper were deposited in zenodo DOI:10.5281/zenodo.5823398. All scripts and data tables for WGCNA analyses are in GitHub https://github.com/lmesrop/WGCNA-bulk-RNAseq-and-single-cell-RNAseq-Hydra-Project.

## Acknowledgements

We thank Rob Steele for Hv105 animals, advice on rearing *Hydra*, and feedback on this manuscript; Stefan Siebert for advice on *Hydra* single cell dissociation; Klebea Carvalho, Gabriela Balderrama-Gutierrez, Katherine Williams, and Camden Jansen for advice on processing and analyzing single cell data.

## Author Contributions

AMM and AM designed the study. AMM and HL conducted experiments. AMM analyzed single cell RNA-seq and bulk ATAC-seq. LYM analyzed bulk RNA-seq. AMM wrote the manuscript with consultation from all authors.

## Supplemental Material

**Figure S1. Bulk RNA-seq expression.** A) MDS plot of bulk RNA-seq samples showing scatter based on gene expression profiles. B) Heatmap of genes highly expressed early in regeneration t0-t4. C) Heatmap of genes upregulated at t12. D) Heatmap of genes upregulated at t48.

**Figure S2. Single cell RNA-seq sample distribution and clustering.** A) Violin plot of nFeatures per sample. B). Violin plot of nCounts per sample. C) UMAP of cell clusters before annotations.

**Figure S3. Clustering and expression of cell marker and candidate regeneration genes.** A) Wnt3. B) KS1. C) Nematogalectin A. D) PPOD1. E) LWamide. F) nematocyte-specific; uncharacterized.

**Figure S4. Expression trajectory in pseudotime for Wnt signaling, early wound response genes, and candidate Wnt3 regulating transcription factors.**

**Figure S5. Slingshot results.** A) Seurat single cell clusters with linages superimposed. B) Range of Seurat clusters expressed in pseudotime. C) Cells colored in pseudotime. D) Gene expression of *Wnt3* in pseudotime. E) Cells aligned in pseudotime trajectory.

**Table S1. Genes upregulated after head bisection**

**Table S2. Genes upregulated at 12 hours post-bisection**

**Table S3. Genes upregulated at 48 hours post-bisection**

**Table S4. Gene list for coral2 module**

**Table S5. Gene list for darkgreen module**

**Table S6. *Cis*-regulation for regeneration genes**

**Table S7. scRNA-seq statistics**

**Table S8. Top markers for seurat clusters**

**Table S9. Dfam IDs for transposable elements**

